# Evolutionary Tree in Chemical Space of Natural Products

**DOI:** 10.1101/2025.08.31.673394

**Authors:** Bo Yang, Chijian Xiang, Tongtong Li, Boyuan Liu, Anton V. Sinitskiy, Jianing Li

## Abstract

Natural products (NPs) are key to biological function and adaptation, with their distribution shaped by complex evolutionary and ecological forces. While it may seem reasonable to assume that closely related species produce chemically similar NPs, this assumption has not been systematically tested at a broad taxonomic scale. Here, we evaluate whether evolutionary (taxonomic) proximity correlates with chemical similarity in large-scale data from the Lotus database of NPs. We use five deep learning-based encoders, including Chemformer and SMILES Transformer, to embed NPs into a high-dimensional “chemical space.” Our results demonstrate that, for flowering plants (Magnoliopsida) and conifers (Pinopsida), species separated by shorter taxonomic distances tend to produce significantly more similar NPs. Similar trends are observed for Fungi and Metazoa, albeit with some complications, possibly due to horizontal gene transfer, convergent evolution, and/or incomplete coverage in the dataset used for NPs. Our findings suggest that the evolutionary tree can be statistically recovered in a chemical space of NPs, provided that this space is constructed with appropriate deep learning techniques, and provide a new computational framework to investigate the evolutionary dynamics of secondary metabolism. These results can inform drug design strategies, for example by enabling the reconstruction of NPs from poorly studied or extinct species.

## Introduction

Closely related species often synthesize chemically similar natural products (NPs). Multiple studies reveal that phylogenetic proximity correlates with shared biosynthetic gene clusters (BGCs) and hence, with structurally related metabolites.^1-26^ For instance, actinomycetes such as *Streptomyces* conserve antibiotic-producing BGCs,^11,12^ and comparable clustering is seen in *Salinispora* and other taxa.^13,16^ Vertical inheritance, gene duplication, and homologous recombination maintain common enzymatic frameworks; modular enzymes such as polyketide synthases and non-ribosomal peptide synthetases diversify products yet preserve core scaffolds within clades.^16^ Comparative metabolomics supports this pattern: metabolite fingerprints in myxobacteria^11^ and the distribution of isoquinoline alkaloids in plants^1^ mirror phylogenetic groupings. The dynamic chemical matrix evolutionary hypothesis proposes that selection for ecological fitness channels related species toward overlapping chemical repertoires.^16,20^

However, many observations contradict a simple one-to-one mapping between evolutionary and chemical proximity. Identical or closely related metabolites, such as nicotine, appear in distantly related plants, and even congeners within the mint family generate divergent terpenoid profiles.^1^ Among bacteria, horizontal gene transfer (HGT), enzyme promiscuity, and rapid mutation remodel BGCs so that neighboring species can produce distinct compounds, such as clavam variants in *Streptomyces clavuligerus* and *S. antibioticus*.^13^ The One Strain–Many Compounds paradigm states that a single genome can yield multiple metabolomes under different conditions, further decoupling chemistry from phylogeny.^16^ Recombination of biosynthetic domains, including non-modular type III polyketide synthases, drives unique pathways even among close relatives.^13,16^ Convergent evolution also generates chemically similar molecules in unrelated lineages. For example, metabolic plasticity in *Streptomyces* responds to environment rather than lineage.^16^ Enzyme promiscuity further contributes to chemical diversity by enabling enzymes to act on diverse substrates, producing a wide range of metabolites.^27^ Together, these factors explain why large-scale surveys report both congruent and incongruent patterns between taxonomy and chemistry.^1-4,6,8-13,15,16,18-22,26,28-31^

Given this mixed evidence, it is critical what tools are used for analysis. To date, most systematic links between evolution and chemistry have traced BGC phylogeny: homologous genes are aligned, phylogenetic trees of catalytic domains are built, and genome-mining tools classify clusters to predict metabolites.^11,17-19^ Comparative genomics reveals duplication, recombination, and horizontal transfer events that shape NP diversity. Phylogenetic profiling shows that metabolite suites are often more similar within clades, guiding prioritization of underexplored taxa such as cyanobacteria and *Streptomyces*.^11,19^ Environmental metagenomes extend discovery to unculturable organisms, and machine learning (ML) models integrate sequencing and chemical data to predict novel products.^18,19^ Yet these efforts typically interrogate individual genera or families; they seldom test broad sections of the evolutionary tree.

In this work, we **hypothesize** that an evolutionary tree can be statistically recovered in a high-dimensional chemical space of NPs. Unlike conventional phylograms where each species is a single node, here every species is represented by a cloud of molecular points scattered through chemical space. Two related species thus correspond to two overlapping point clouds rather than adjacent nodes, complicating visualization and demanding quantitative metrics. Our strategy is to compute chemical distances between sets of NP from the Lotus database^32,33^ for various pairs of biological species, and then, use standard statistical tests to check whether *chemical distances for smaller taxonomic distances* are statistically significantly *smaller* than *chemical distances for greater taxonomic distances*. In brief, this formalization of our hypothesis can be worded as “**Closer taxonomy, closer chemistry**”.

It is easy to define **taxonomic distance**, which we use as a proxy for evolutionary distance. The same species has a taxonomic distance of 0 to itself. Two different species within the same genus have a taxonomic distance of 1. If two organisms are from different genera within the same family, they have a taxonomic distance of 2, etc. (We accept the taxonomic schema used by the Lotus dataset.) As for the **chemical distance**, we define it below (see Methods), taking into consideration the above-mentioned factors that complicate the relationship between evolutionary and chemical closeness (HGT, convergent evolution, etc), as well as the incompleteness of existing NP databases.

## Results

### Chemical distances at shorter taxonomic distances are significantly smaller than those at greater taxonomic distances

Specifically, NPs from species within the same genus (taxonomic distance of 1) exhibit significantly lower chemical distances than those from species in different genera within the same family (taxonomic distance of 2). Similar relationships are observed when comparing species from different genera within the same family (taxonomic distance of 2) or species within the same genus (taxonomic distance of 1) to species from different families within the same class (taxonomic distance of 3).

To statistically validate this pattern, we applied Welch’s t-test to compare the distributions of chemical distances across taxonomic distances. A one-tailed test was used to assess whether chemical distances *increase* as species become more distantly related. The results reveal low p-values for the comparisons between taxonomic distances 1 and 2 (*p* = 3.5·10^−30^) and between taxonomic distances 1 and 3 (7.6·10^−105^), indicating highly statistically significant differences, in strong support of the hypothesis of this paper (Table 1, row “All”). As for the comparison of chemical distances between taxonomic distances 2 and 3, the resulting p-value is exceptionally low (4.3·10^−448^), which we explain by an order of magnitude greater effective number of degrees of freedom for this comparison than for the first two comparisons, which, in turn, can be easily explained by the branching nature of the evolutionary tree.

**Table 1.** Statistical significance (p-values) and effective number of degrees of freedom (e.d.o.f.) for comparisons of chemical distances at different taxonomic distances. P-values less than 0.01 are shown in bold. For the definition of chemical distances, see Methods.

| Part of the evolutionary tree | Taxonomic distances |  |  |  |  |  |
| --- | --- | --- | --- | --- | --- | --- |
|  | 1 vs 2 |  | 2 vs 3 |  | 1 vs 3 |  |
|  | p-value | e.d.o.f. | p-value | e.d.o.f. | p-value | e.d.o.f. |
| All | **$3.5 \cdot 10^{-30}$** | 503. | **$4.3 \cdot 10^{-448}$** | 4016. | **$7.6 \cdot 10^{-105}$** | 417. |
| Kingdom Viridiplantae | **$4.6 \cdot 10^{-34}$** | 275. | **$5.2 \cdot 10^{-425}$** | 3803. | **$1.9 \cdot 10^{-75}$** | 246. |
| Class Magnoliopsida (flowering plants) | **$5.1 \cdot 10^{-29}$** | 249. | **$7.4 \cdot 10^{-410}$** | 3724. | **$6.5 \cdot 10^{-66}$** | 226. |
| Class Pinopsida (conifers) | **$6.3 \cdot 10^{-13}$** | 54. | **$3.1 \cdot 10^{-10}$** | 96. | **$2.6 \cdot 10^{-17}$** | 27. |
| Kingdom Fungi | 0.014 | 289. | **0.0082** | 450. | **$5.7 \cdot 10^{-6}$** | 335. |
| Kingdom Metazoa (animals) | 0.026 | 28. | 0.17 | 19. | **$7.9 \cdot 10^{-7}$** | 29. |

In this analysis, each computed interspecies chemical distance received equal weighting, provided it could be reliably calculated from the Lotus dataset (see Methods). In fact, the majority of reliable chemical distances in the Lotus dataset originated from flowering plants, which were well-represented with 44 reference species across 27 families. In contrast, conifers (5 species across 2 families), fungi (9 species across 6 families) and animals (9 species across 8 families) had poorer representation. To address these discrepancies in taxonomic coverage, we repeated our analyses separately within each of these major taxonomic groups.

### Statistical confidence of these conclusions, given the Lotus dataset used, varies across taxonomic groups

The evidence supporting the hypothesis—that chemical distances increase with greater evolutionary distance—is very strong for plants (Magnoliopsida and Pinopsida) but becomes weaker or absent as we move to fungi and animals (Table 1). Thus, the support for this hypothesis from the Lotus data depends on the specific region of the evolutionary tree analyzed. The overall statistics are primarily driven by flowering plants which have both extensive representation in the dataset and highly significant statistical support. On the other hand, NPs from animals have a poorer representation in this database. In particular, the only p-value much greater than 0.01 in Table 1 is observed for a comparison of taxonomic distances of 2 and 3 in animals. Note that the effective number of degrees of freedom in this case is more than two orders of magnitude smaller than that of flowering plants. In addition to insufficient data, higher p-values may be caused by HGT or convergent evolution, as discussed in the Introduction.

Next, we will refine our analysis by examining distributions of p-values, each of which is computed for a certain reference species. This approach will provide deeper insights into the consistency of observed trends across species and help identify potential variability and its causes within each taxonomic group. We combine this analysis with consideration of how these distributions of p-values are affected by the choices of (1) the encoders defining the chemical space, and (2) the parameters involved in the definition of the chemical distance.

### Various deep learning-based molecular encoders can construct a “chemical space” that enables the mapping of evolutionary relationships through chemical similarity

For each embedding, we computed p-values for each reference species and built statistical distributions of the resulting p-values. Then, we compared the resulting distributions for different embeddings (Fig 1). Comparisons were made for taxonomic distances of 1 and 3, to clarify how the lowest p-values from the meta-analysis across distant taxonomic groups (Table 1) are manifested at the level of specific species.

**Fig. 1.**
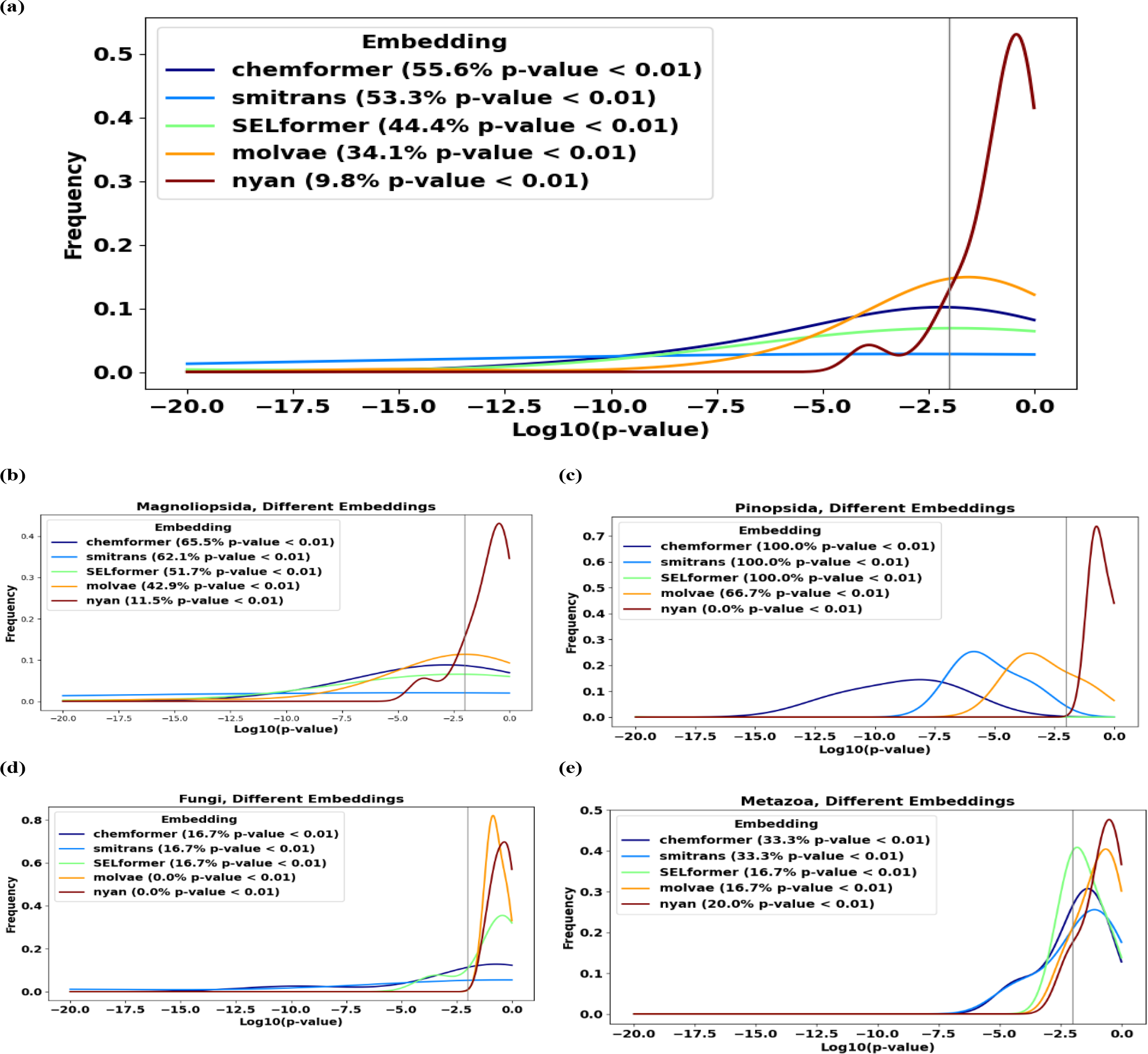
(a) Chemformer and SMILES Transformer (“smitrans” in figure legends) embeddings can build chemical spaces of natural products, such that the hypothesis “Closer taxonomy, closer chemistry” is statistically confirmed (p-values < 0.01) not only at the meta-level, but also at the level of individual reference species in more than half of such species available from the Lotus dataset. SELFormer and Molecular VAE (“molvae” in figure legends) embeddings demonstrate poorer performance in this task, and the performance of the NYAN embedding is inferior. The fractions of reference species with the confirmed hypothesis depend on the branches of the evolutionary tree: they are higher in (b) flowering plants and (c) conifers than in (d) fungi and (e) animals.

Among the five tested embeddings, Chemformer demonstrated the strongest and most consistent performance (Fig. 1, *dark-blue curves*). Even though the statistics at the resolution level of individual reference species are limited, for more than half (namely, 55.6%) of all reference species, the log_10_ p-values remained below -2 (that is, *p* < 0.01) with this encoding, with some cases reaching values as strong as log_10_ p-values < -20 (*Artemisia annua, Isodon rubescens*, both flowering plants). This indicates strong statistical support for the relationship between chemical and evolutionary distances even at the level of individual reference species.

SMILES Transformer and, to a lesser degree, SELFormer (Fig. 1, *light-blue and green curves*) also exhibited robust performance, with log_10_ p-values < -2 for nearly half of all reference species, though less than with Chemformer. In contrast, Molecular VAE and NYAN embeddings (*orange and dark-red curves*) performed significantly worse, often yielding statistically insignificant p-values >> 0.01.

Across four main taxonomic groups of reference species in the available data (Magnoliopsida, Pinopsida, Fungi, Metazoa), the relative performance of the encoders stayed consistent, with Chemformer and SMILES Transformer demonstrating the highest performance. At the level of individual reference species, statistically significant p-values were observed in all conifers and approximately 2/3 of flowering plants, while in fungi and animals this fraction was lower (Fig. 1b-e). The latter observation can be at least partially explained by poorer statistics at the level of individual species, requiring a meta-analysis at higher-level taxonomic groups, such as kingdoms, to reveal statistically significant support of the hypothesis, as performed in the previous subsections in the Results.

Despite these limitations, it is impressive that taxonomic relationships between biological species, genera and families can be statistically significantly detected in a chemical space of NPs, *even though the chemical space is built with a deep learning embedding trained for entirely different tasks* without using any evolutionary data, and moreover, *for various deep learning models* (in this work, Chemformer, SMILES Transformer and, to a lesser degree, SELFormer) trained to meet different goals.

### Chemformer embedding not only provides strong statistical confirmation of the hypothesis but also remains highly robust across a wide range of hyperparameter settings

To evaluate this robustness, we examined the impact of two key parameters: (i) the minimum number of NPs required per species (threshold size) to be included in the analysis as a current species, and (ii) the percentile used to define chemical distances. Across a wide range of their values, the fraction of reference species with p-values below 0.01 remained nearly unchanged (Fig. 2).

**Fig. 2.**
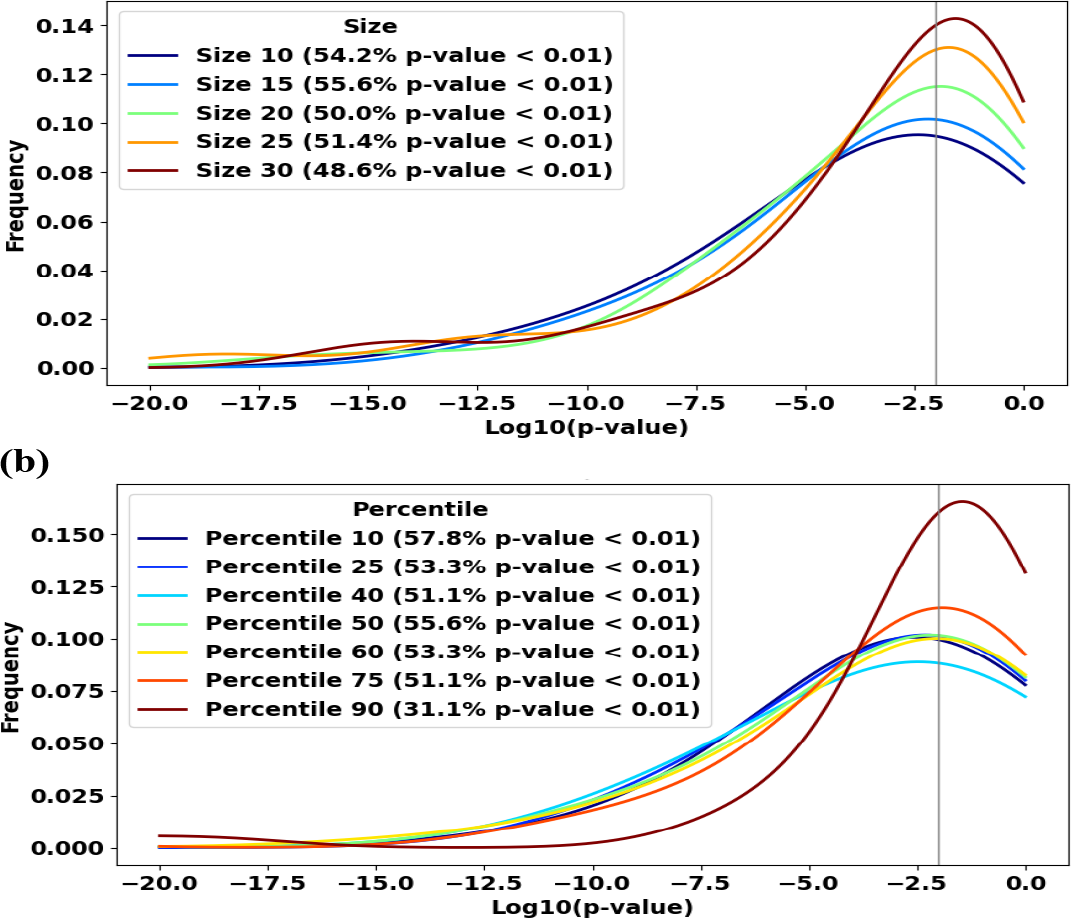
Varying (a) the threshold size in a broad range of reasonable values, namely 10 to 30, and (b) the percentile levels in a broad range of reasonable values, namely 25 to 75, do not strongly affect the distributions of p-values for reference species. Chemformer was used to build the chemical space of natural products.

The default value for the threshold size, namely 15, ensures the maximum fraction of reference species with *p* < 0.01. However, this maximum is shallow and wide, with sizes down to 10 and up to 30, resulting in decreases in this fraction by only a few percent (Fig. 2a). Note that the fractions of species within Pinopsida and Fungi do not change in the wide range of threshold sizes from 10 to 30. Within Metazoa, the performance is better for larger sizes (25 and 30, in comparison to 10, 15 and 20), but this effect is overpowered by the maximum performance within Magnoliopsida at the threshold size of 15. Overall, in the broad range of the threshold size from 10 to 30, the sensitivity of the fraction of reference species with *p* < 0.01 to this parameter is much weaker than to the choices of the encoder or the high-level taxonomic group for which the performance is evaluated.

The default value for the percentile level, namely 50, ensures a local maximum fraction of reference species with *p* < 0.01. This maximum is also shallow and wide, with percentile levels in a reasonable range of 25 to 75 providing decreases in this fraction by only a few percent (Fig. 2b). The fraction of reference species with *p* < 0.01 significantly decreases only at the unreasonably high level of this parameter, namely 90, which is expected to make interspecies chemical distances too sensitive to missing NPs in the database. As for the percentile level of 10, it provides a better performance than the default value of 50 that we choose. However, this low value of the percentile level would likewise make interspecies chemical distances too sensitive to incomplete data, given the small optimal threshold size. From the perspective of high-level taxonomic groups, in Pinopsida and Fungi, the fractions of species with *p* < 0.01 do not change in the wide range of percentile levels except for the unreasonably high value of 90, while both in Magnoliopsida and Metazoa, they reach local maxima at the percentile level of 50. Similarly to the previous parameter, the sensitivity of the fraction of reference species with *p* < 0.01 to the percentile level, in the reasonable range of its values from 25 to 75, is overall much weaker than the sensitivity to the choices of the encoder or the large taxonomic group.

### Visualization of typical examples of pairs of natural products separated by various taxonomic distances illustrates the conclusion that evolutionarily closer species have closer chemical structures of natural products

Consider two typical examples.

#### Example 1

*Tripterygium wilfordii* (“thunder god vine”, a plant used in traditional Chinese medicine) happens to be the reference species with the biggest number of NPs in the Lotus dataset (488 NPs). At **taxonomic distance of 1** from *T. wilfordii*, only two species in the Lotus dataset (*T. hypoglaucum* and *T. doianum*) have at least 15 NPs (the threshold level introduced above). In these two pairs (Fig. 3a), scaffolds of the molecules are the same (taking into account the keto-enol tautomerization in the first pair), and only some of the functional groups differ (e.g., -CH_3_ vs -CHO, >C=O vs -CH_2_-, etc.). At **taxonomic distance of 2**, 35 species in the Lotus dataset have at least 15 NPs. Among them, typical (closest to the average) distances in the chemical space are observed for the molecular pairs shown in Fig. 3b. In these pairs, scaffolds of the molecules are similar, but not exactly the same. At **taxonomic distance of 3**, as many as 2795 species in the Lotus dataset have at least 15 NPs. For typical (closest to the average) distances in the chemical space (Fig. 3c), in these pairs, unlike those for the taxonomic distances of 1 or 2, scaffolds of the molecules are essentially different, including different numbers and sizes of rings and different heteroatoms involved in them. Thus, it is evident even with a naked eye that dissimilarity of the molecules in the pairs increases with the taxonomic distance from *T. wilfordii*. This visualization confirms the numerical statistical conclusions given in the previous subsections.

**Fig. 3.**
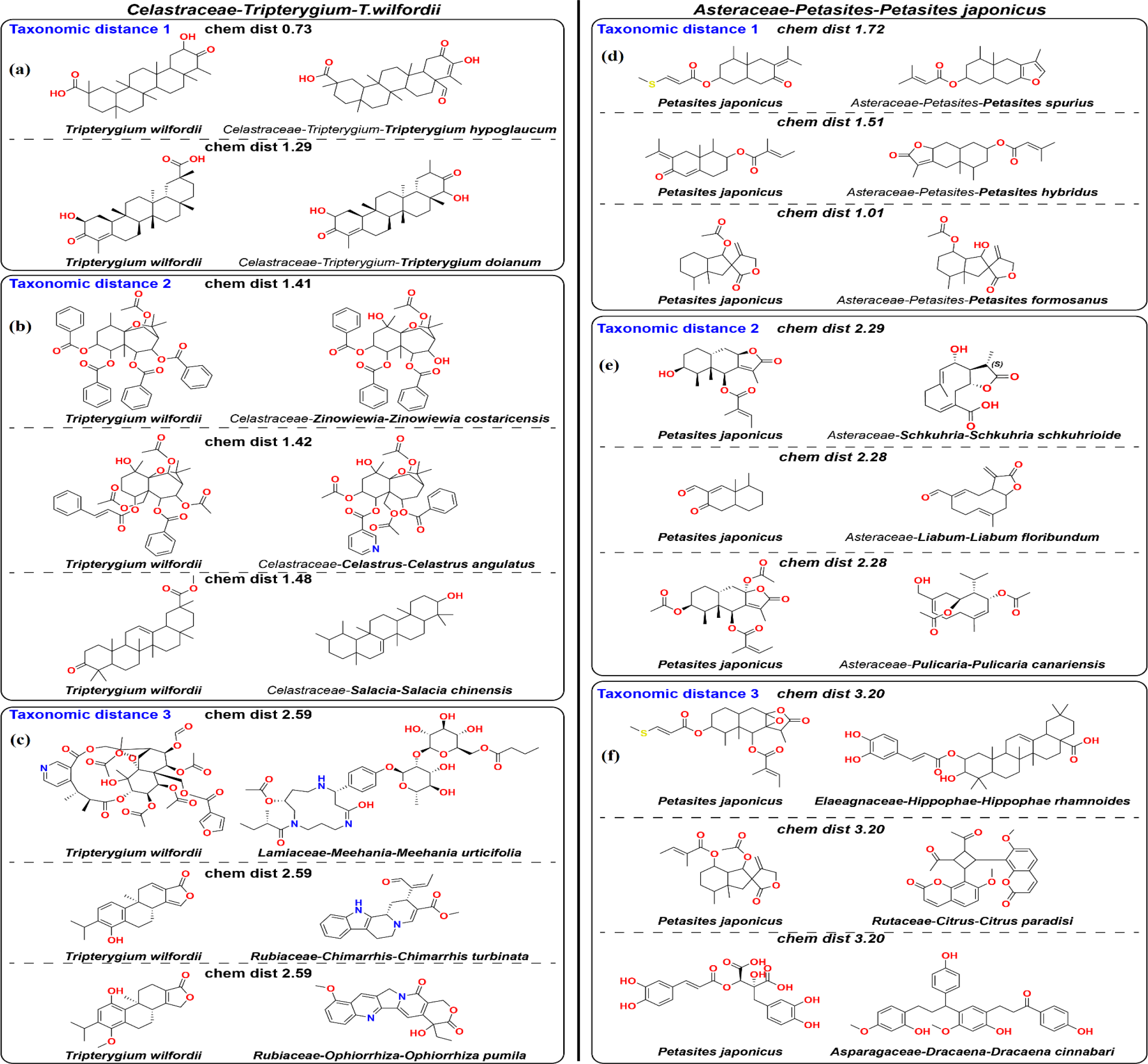
Visualization of typical pairs of molecules corresponding to taxonomic distances of 1, 2, and 3 demonstrates that, in typical cases, the hypothesis “Closer taxonomy, closer chemistry” is supported. In all pairs, a structure on the right is a natural product from a current species (mentioned below the structure), and a structure on the left is the closest natural product from the reference species (*Tripterygium wilfordii* in the left panels and *Petasites japonicus* in the right panels).

To additionally illustrate these patterns, we projected the high-dimensional chemical space onto two dimensions using tSNE (Fig. 4). Despite this reduction, evolutionary structure remains hard to visualize, as each species is represented by many points (each corresponding to one NP). In Fig. 4a, points are colored by taxonomic distance from *T. wilfordii*. Further simplifying this, Fig. 4b selects a single NP from *T. wilfordii* and plots only the nearest NP from each species. This figure illustrates the trend that median chemical distances increase with taxonomic distance, a pattern previously demonstrated statistically across the whole dataset.

**Fig. 4.**
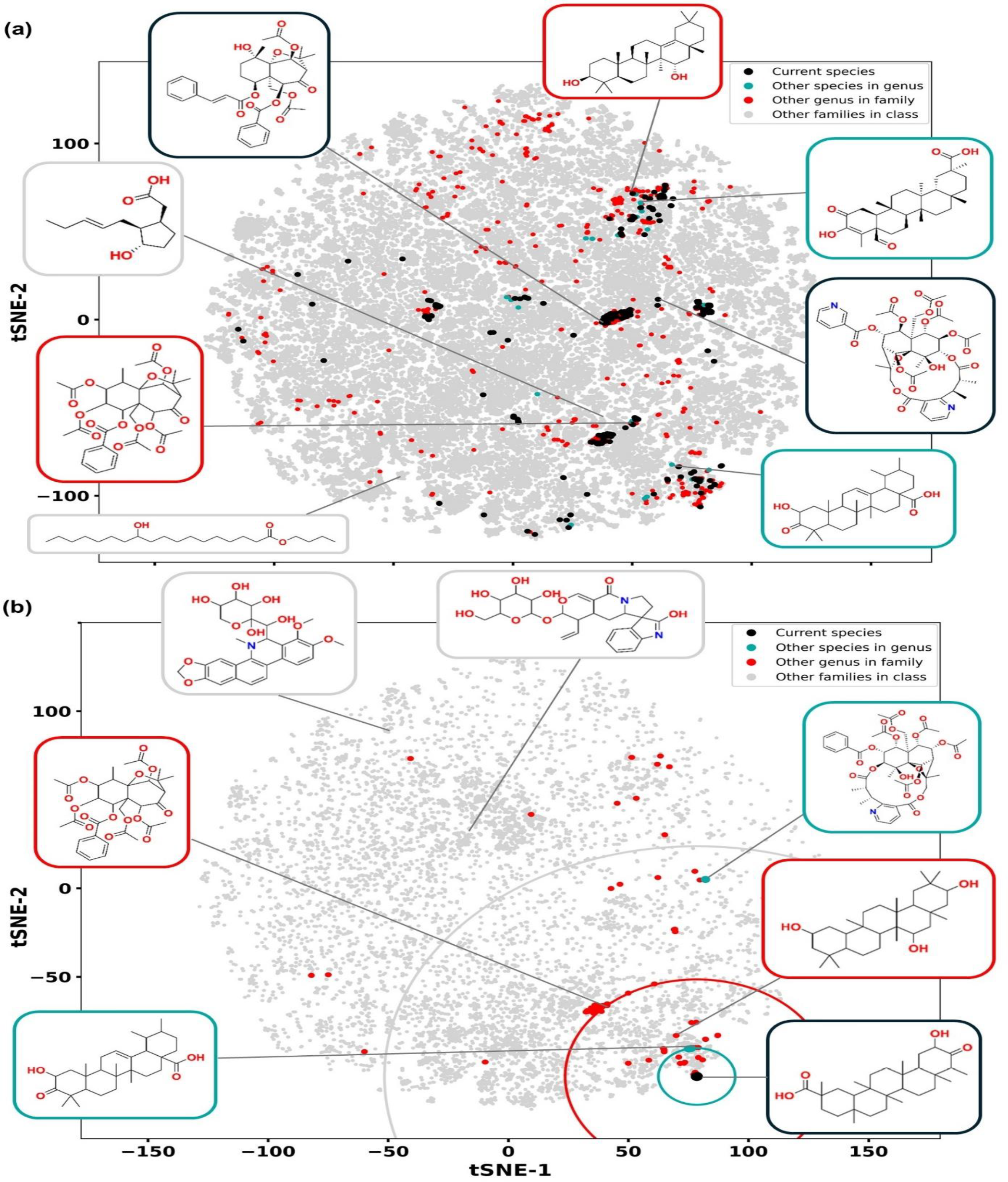
Visualizing evolutionary relationships in NP chemical space. (a) NPs from a given species (here, *T. wilfordii*, shown as black dots in a 2D tSNE projection of all flowering plant NPs) are dispersed across distant regions of chemical space. They tend to be surrounded by NPs from species with closer taxonomic distances (*cyan*, t.d. 1; *red*, t.d. 2; *grey*, t.d. 3), though this reflects a statistical trend rather than a strict rule. (b) For a selected NP (black dot, a structure shown in Fig. 3a), we plot one NP per species corresponding to the closest match in chemical space. NPs from more closely related species (*cyan* and *red*) tend to lie nearer than those from more distant species (*grey*). Colored circles indicate median chemical distances for each taxonomic distance.

#### Example 2

Among the p-values, the distributions of which are plotted in Fig. 1 (for Chemformer) and Fig. 2 (for size 15 in panel (a) and percentile 50 in panel (b)), the median value (namely, *p* = 0.0077) is achieved for *Petasites japonicus* (family *Asteraceae*) as a reference species. This is another flowering plant, also known as “butterbur”, and having 155 NPs in the Lotus dataset. At **taxonomic distance of 1** from *P. japonicus*, only four species in the Lotus dataset have at least 15 NPs. Typical (closest to the average) distances in the chemical space correspond to the pairs of molecules shown in Fig. 3d. In the first two pairs, the scaffolds are different, but the -C(=O)-CR=CR-fragment in the structures from *P. japonicus* are conformationally rigid and close in geometry and partial charge distribution to the flat furan rings in the corresponding structures on the right hand side. As for the third pair, the scaffolds there are the same, and functional groups are similar. At **taxonomic distance of 2**, 530 species in the Lotus dataset have at least 15 NPs. Typical (closest to the average) distances in the chemical space are shown in Fig. 3e. At first sight, the scaffolds in all three pairs are significantly different. However, note that the 10-membered (cyclodecadiene) rings on the right and fused 6-membered (decalin) rings on the left differ by a formation of a single C-C bond to form the fused ring system, similar to structures involved in biochemical reactions catalyzed by terpene synthases. At **taxonomic distance of 3**, 2298 species in the Lotus dataset have at least 15 NPs. In the pairs of molecules with the typical chemical distances (Fig. 3f), the differences between scaffolds are significantly greater, and such transitions could not be simply performed with a small number of biochemical reactions. Overall, the case of *P. japonicus* also provides a clear visual illustration of the trend of increasing chemical dissimilarity with the increase in the taxonomic distance, going from similar or identical scaffolds to different scaffolds related by possible biochemical reactions, and then to scaffolds very dissimilar biochemically. Despite the fact that the p-value for this reference species, *p* = 0.0077, is close to the conventional threshold for statistical significance, *p* = 0.01, a clear distinction takes place between chemical similarities of NPs at taxonomic distances of 1, 2, and 3.

Additionally, comparing these two examples demonstrates that in different branches of the evolutionary tree, varying degrees of chemical similarity may correspond to taxonomic distances of 1 and 2 (more similarity in each case for *T. wilfordii* and less similarity in the corresponding cases for *P. japonicus*). This suggests possible limitations of the analysis provided in Table 1.

### Reference species in Fungi and Metazoa with high p-values may be more prone to horizontal gene transfer

Among the high-level taxonomic groups, Fungi and Metazoa have the lowest fraction of reference species with *p* < 0.01 (Fig. 1). To clarify possible reasons for this, we consider these cases in more detail.

With the default settings, six reference species from Fungi have enough data in the Lotus dataset. In five of them (*Alternaria alternata, Aspergillus fumigatus, Aspergillus ustus, Penicillium chrysogenum, Penicillium citrinum*), p-values are statistically insignificant (ranging from 0.12 to 0.47), while for one of them, *Ganoderma applanatum*, statistical significance is very strong (*p* = 1.7·10^-10^). It is noteworthy that the first five species are microfungi and belong to phylum *Ascomycota*, while the last one is macroscopic and belongs to phylum *Basidiomycota*. A plausible explanation might be that HGT is generally more significant in microfungi due to ecological, genomic, and lifestyle factors.^34-36^ However, the data available from the Lotus dataset are not sufficient to provide a final answer to this question. Further research is required to establish the degree of certainty regarding the hypothesis “Closer taxonomy, closer chemistry” in the evolutionary branch of microfungi/Ascomycota.

Among Metazoa, six reference species have sufficient data in the Lotus dataset. In four of them (*Clavularia viridis, Aplysia dactylomela, Sinularia flexibilis, Briareum excavatum*), p-values are statistically insignificant (ranging from 0.016 to 0.15), while for the remaining two (*Briareum asbestinum, Antillogorgia elisabethae*), statistical significance is strong (*p* = 5.0·10^-5^ and 0.0040, respectively). Interestingly, the latter two species, unlike the former four, are Caribbean species,^37^ which might create less favorable ecological conditions for HGT. As in the case of fungal species listed above, further research is required to assess the applicability of the hypothesis “Closer taxonomy, closer chemistry” in the evolutionary branch of Metazoa, and more specifically, corals and mollusks, to which these six species belong.

## Discussion

Our analysis supports the hypothesis that the evolutionary tree is partially reflected in the chemical space of NPs. We show that chemical distances among species correlate with taxonomic distances, a pattern statistically robust across multiple deep-learning-based molecular embeddings. However, the strength of this relationship varies across taxonomic groups, shaped by the biological complexity of NP biosynthesis and the limitations of the Lotus dataset.

The most consistent correlation is found in plants. This aligns with prior work showing phylogenetic clustering of BGCs and structural conservation of metabolites within related clades. Our results extend these findings across broader taxonomic scales, revealing that species within the same genus tend to produce more chemically similar NPs than those from different genera or families. This may be due to the relative conservation of biosynthetic pathways in plants, along with coevolutionary pressures such as interactions with herbivores, microbes, and pollinators. These results suggest that in plants, vertical inheritance, gene duplication, and gradual divergence of biosynthetic machinery largely govern chemical diversity.

In contrast, fungi and animals (Metazoa) show weaker correlations between chemical and evolutionary distances. This likely reflects greater influences of HGT, convergent evolution, and metabolic plasticity in these groups. In fungi and animals, BGCs are more readily exchanged across unrelated taxa, leading to chemically similar compounds arising in distantly related species. Moreover, environmental factors often drive metabolic responses, further decoupling NP profiles from evolutionary relatedness. These complexities highlight the need for caution when interpreting phylogenetic signals in NP chemistry.

These findings have practical implications for NP discovery. In groups where chemical and evolutionary proximity align, phylogenetic information can guide bioprospecting efforts. Closely related species within well-characterized clades are more likely to yield structurally related bioactive compounds. In contrast, for fungi and animals, approaches that incorporate genomic or ecological data may be more effective, given the prevalence of HGT and adaptive metabolic diversity. From an evolutionary perspective, our results reinforce the idea that NP biosynthesis evolves under both genetic and ecological constraints. The modular nature of biosynthetic enzymes, selective pressures, and stochastic evolutionary events together shape chemical diversity across lineages. Our approach, which integrates machine learning and large-scale chemical databases, offers a quantitative framework for examining these patterns on a global scale.

Limitations of this work include the following. NP databases, including the Lotus dataset used here, are incomplete, particularly for underexplored taxa. Broader coverage would improve resolution and confidence in future analyses. Additionally, chemical space is multidimensional and sensitive to the choice of embedding model. While Chemformer performed best in our study, further development of models tailored to evolutionary analysis may yield improved results.

In conclusion, this study shows that taxonomic signals can be detected in the chemical space of NPs, particularly in plants. Deep-learning tools provide a promising route for investigating the evolutionary dynamics of natural product biosynthesis across the tree of life. Moving forward, integrating metabolomics, genomics, and machine learning will be essential for deepening our understanding of the complex relationship between evolution and chemistry of NPs. These insights can inform rational drug discovery by enabling the prediction or reconstruction of NPs from poorly characterized or extinct lineages. Leveraging evolutionary proximity in well-mapped clades may help identify structurally novel bioactive compounds that are otherwise inaccessible due to gaps in empirical NP data.

## Methods

### Encoders to build chemical space of natural products

To construct a chemically meaningful space for NPs, we used deep learning-based molecular encoders to generate high-dimensional vector representations of molecular structures (Fig. 5a). These embeddings enable quantification of chemical similarity between species based on their NP compositions. We evaluated five encoders: Chemformer,^38,39^ SMILES Transformer,^40,41^ SELFormer,^42,43^ NotYetAnotherNightshade (NYAN),^44,45^ and the historically first chemical VAE.^46^ For the latter, we used a PyTorch implementation^47^ with a pretrained model; to avoid confusion with the original model, we will refer to it as “molvae”.

**Fig. 5.**
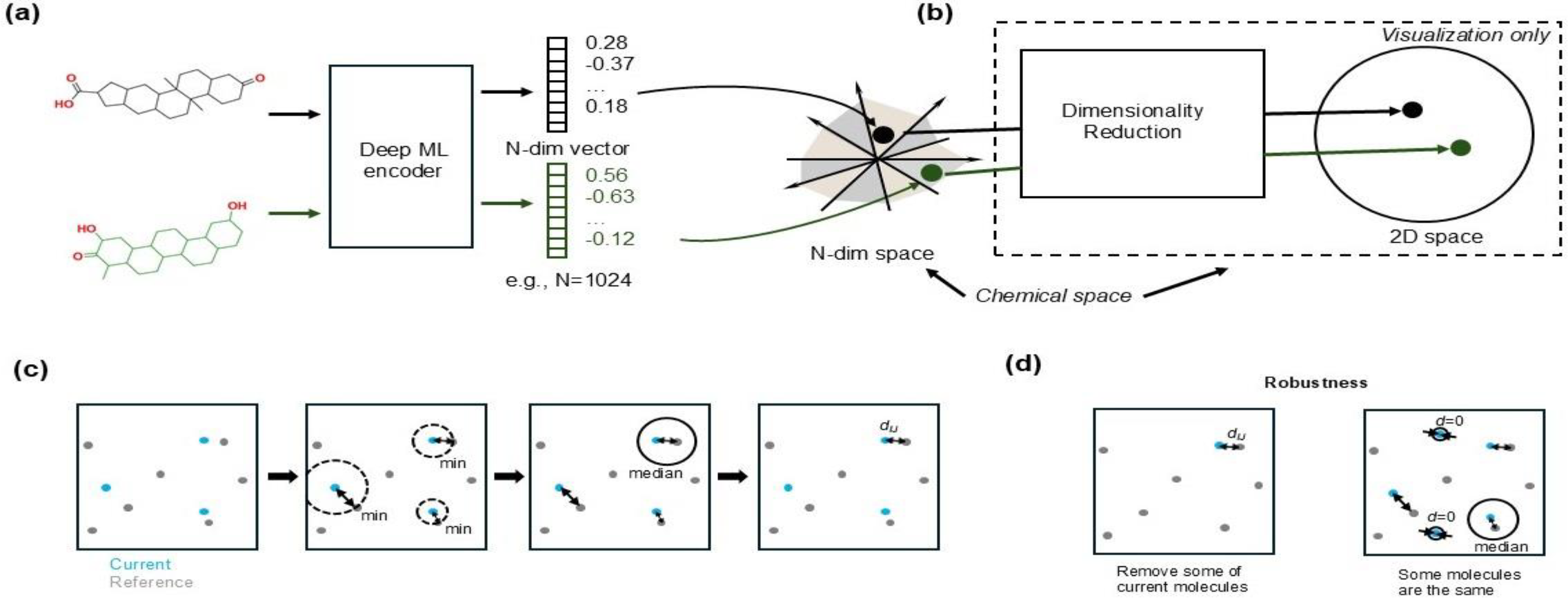
Construction and interpretation of chemical space and interspecies distances using deep learning-based molecular embeddings. (a) Deep learning models project NP chemical structures onto a high-dimensional Euclidean space. (b) For visualization, tSNE is used to reduce this space to two dimensions. (c) Chemical distance between NP sets is defined as follows. Structures are embedded in high-dimensional space. For each NP in the current species (*blue*), the closest NP in the reference species (*gray*) is identified. The median of these distances is the species-to-species distance. (d) This approach is robust to missing data and ubiquitous metabolites.

None of the models were retrained or provided with evolutionary data during training. For visualization (Figs. 4, 5b), we projected the chemical space onto two dimensions using tSNE with default parameters; however, all chemical distances were computed in the original high-dimensional space.

### Definition of chemical distance between species

We define the chemical distance between two biological species as the dissimilarity between their respective sets of NPs, computed from the molecular embeddings (Fig. 5c). The calculation proceeds in three steps:

1. Molecular Distance: The chemical distance *d*_*nm*_ between two molecules (NPs) *n* and *m* is defined as the Euclidean distance between their embeddings in a high-dimensional chemical space.
2. Molecule-to-Species Distance: For a molecule *n* in a current species *I*, we calculate the minimum distance *d*_*nJ*_ to any molecule *m* in a reference species *J*:

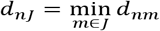

We assume that the number of molecules known for the reference species is large relative to that of other species in the used dataset. By construction, the list of known NPs from the reference species is more complete than for the current species, therefore, this sort of minimization (unlike a minimization over all molecules in the less well-studied current species) will provide a more robust estimate of the distance aggregated to the level of a species.
3. Species-to-Species Distance: For species *I* and *J*, we define the overall chemical distance *d*_*IJ*_ as the percentile *P*_*k*_ of the set {*d*_*nJ*_}, where *n* runs over all molecules in species *I*:

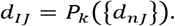

We use the median (50th percentile) by default. Percentile-based aggregation helps correct for biases from missing or overly common NPs. For instance, unusually high *d*_*nJ*_ values may reflect missing similar NPs in the reference species, while unusually low values may correspond to universally shared metabolites (Fig. 5d). Percentile selection (0<*k*<100) ensures robustness and interpretability. This definition of chemical distance is asymmetric, depending on which species is taken as the reference, a necessary choice given the imbalance of the Lotus dataset.

### Parameters in the definition of chemical distance

The chemical distance *d*_*IJ*_ depends on the following key parameters:

- Encoder model used to construct molecular embeddings.
- Percentile level *k* used to compute *d*_*IJ*_ from the *d*_*nJ*_ values.

Additional parameters apply when aggregating data across species in statistical analysis. To ensure robust estimates, we exclude species with few reported NPs. Thus, we set thresholds for inclusion:

- Minimum number of NPs per species to serve as a current species.
- Minimum number of NPs per species to serve as a reference species.

More than half the species in the Lotus dataset have six or fewer NPs, so this filtering is critical to minimize statistical artifacts. Unless otherwise noted, we accept the following default values of these parameters in this work: the encoder is Chemformer, the percentage *k* is 50 (i.e., *d*_*IJ*_ is the median value of {*d*_*nJ*_}), and the threshold levels for the number of NPs are 15 and 150 for inclusion as current and reference species, respectively. These default values provide the strongest and most stable statistical support for our hypothesis “Closer taxonomy, closer chemistry,” given the data from the Lotus dataset.

## Acknowledgement

We thank Prof. Ray Bressan, Brian Metzger and Zhong-Yin Zhang (Purdue University) for helpful discussions. This work was supported by an NIH R01 award (R01GM143370) and the AnalytiXIN fellowship.

## Notes

### Competing Interest Statement

The authors have declared no competing interest.

